# *Streptomyces* sp. N2A promotes tomato (*Solanum lycopersicum* L.) vegetative growth and yield by modifying fruit morphology

**DOI:** 10.64898/2026.07.10.737781

**Authors:** Rodrigo Maldonado, Oriana Iacomozzi, Gustavo Rodríguez, Eduardo Rodríguez, María Amalia Chiesa

**Affiliations:** Laboratorio de Eco-Fisiología Vegetal (LEFIVE), Instituto de Investigaciones en Ciencias Agrarias de Rosario, Consejo Nacional de Investigaciones Científicas y Técnicas (IICAR-CONICET), Facultad de Ciencias Agrarias, Universidad Nacional de Rosario (UNR), Zavalla, Santa Fe, Argentina; Grupo de Genética y Mejoramiento de Tomate (GMT), Instituto de Investigaciones en Ciencias Agrarias de Rosario, Consejo Nacional de Investigaciones Científicas y Técnicas (IICAR-CONICET), Facultad de Ciencias Agrarias, Universidad Nacional de Rosario (UNR), Zavalla, Santa Fe, Argentina; Departamento de Microbiología, Instituto de Biología Molecular y Celular de Rosario, Consejo Nacional de Investigaciones Científicas y Técnicas (IBR-CONICET), Facultad de Ciencias Bioquímicas y Farmacéuticas, Universidad Nacional de Rosario (UNR), Ocampo y Esmeralda, 2000 Rosario, Argentina

**Keywords:** Actinobacteria, Biostimulant, Crop-yield, Phytohormones, Plant growth promoting *Streptomyces* (PGPS)

## Abstract

Tomato production, yield and fruit quality face major challenges due to several factors, including the complex polygenic inheritance of agronomically relevant traits, biotic and abiotic stresses, and increasingly stringent regulations limiting the use of phytosanitary products. In this context, bioinoculants have emerged as a sustainable strategy capable of enhancing yield without compromising fruit quality, conferring protection against different stresses and exerting a minimal or no impact on environment and human health. In this study, we evaluated the effects and the underlying mechanisms by which *Streptomyces* sp. N2A, an actinobacteria isolated from soybean rhizosphere, promotes seed germination, vegetative growth and yield in tomato, without modifying fruit quality. The obtained results demonstrated that the bacterial treatment significantly improved seedlinǵs emergence and growth and development in vegetative stage. At harvest, yield was also significantly enhanced, mainly driven by increased individual fruit weight, which was positively correlated with a thicker pericarp in fruits from N2A-treated plants. Transcriptional analysis during fruit development revealed a coordinated induction of auxin and cytokinin signaling pathways before and after anthesis, providing a hormonal framework that underlies the promotion of pericarp growth. This study provides evidence of the beneficial effect of inoculation with *Streptomyces* sp. N2A on tomato yield and constitutes the first report describing the modification of fruit morphology and expression of genes involved in phytohormonal modulation during early growth and development, induced by a plant growth-promoting *Streptomyces*.

## 1. INTRODUCTION

Tomato [*Solanum lycopersicum* L.] is one of the most economically important horticultural crops, with a production of more than 190 million tons worldwide (Food and Agriculture Organization, 2025). Its high level of consumption and global relevance are largely attributed to its nutritional value, particularly rich in vitamins A and C, as well as bioactive compounds such as lycopene and carotenoids, which are widely recognized for their antioxidant properties (Ali et al, 2021). This high production reflects its global importance for both fresh consumption and industrial processing. However, despite the increase in total production over recent decades, achieving further yield gains remains challenging because it is a complex quantitative trait and because of increasingly stringent regulations on the use of phytosanitary products (Lankinen et al, 2024); therefore, additional yield increment will need to be achieved through sustainable strategies that consider the potential environmental impacts of different practices (Bhandari et al, 2023; Food and Agriculture Organization, 2025). Among these, the use of plant growth-promoting rhizobacteria (PGPR) has gained considerable attention due to their ability to enhance nutrient acquisition, modulate phytohormone homeostasis, and promote plant growth and development through multiple direct and indirect mechanisms (de Andrade et al, 2023; Yang et al, 2024). Particularly for tomato crops, it is critical to select and evaluate PGPR capable of not only increasing yield but also ensuring crop health and maintaining fruit quality under an intensified production system (Drobek et al. 2019). Effective PGPR can contribute to disease suppression and improved plant fitness under adverse environmental conditions, supporting sustainable yield increases (Zhang et al, 2024).

Actinobacteria, such as *Streptomyces* spp., produce more than half of known natural products, including a wide range of antibacterial and antifungal metabolites that are widely used in human medicine and others that play an important role in the biocontrol of agriculturally relevant diseases (Vurukonda et al, 2018; Mesquita et al, 2025). In addition to exerting direct antagonistic effects through the production of secondary metabolites, *Streptomyces* spp. can enhance plant resistance by priming host immune responses and activating systemic defense mechanisms before pathogen attack (Kim et al, 2024; Villafañe et al, 2025). These mechanisms, including the phytohormone and siderophore production, enhance resilience to biotic stress and increase tolerance to adverse environmental conditions (Wang et al, 2025).

Despite their considerable potential, only a limited number of commercial bioinoculants based on *Streptomyces* spp. are currently available worldwide, including Mycostop® or Actinovate®. In this sense, *Streptomyces sp.* N2A (N2A) was recently isolated from soybean roots and identified as an endophytic bacterium capable of colonizing internal tissue, enhancing soybean crop yield and phytosanitary quality of seeds, and reducing both, biotic and abiotic stresses impact in soybean plants (Villafañe et al, 2024; Villafañe et al, 2025; Maldonado et al, 2026).

To explore the potential application of *Streptomyces sp.* N2A in horticultural crop systems, in this work we first evaluate the capacity of this strain to promote vegetative growth and development from seed germination through vegetative stage and then assessed its effect on tomato fruit yield and quality. Our findings support the potential use of *Streptomyces sp.* N2A as a biostimulant for horticultural crops and its contribution to more sustainable production systems.

## 2. MATHERIALS AND METHODS

### 2.1 Bacterial strain and culture conditions

*Streptomyces sp*. N2A was grown in International *Streptomyces* Project-2 (ISP2) agar media for spore preparation, and the collected spores were stored at -80°C. Bacterial cultures grown in ISP2 liquid medium at 30°C for two days were used to inoculate tomato seeds with a mixture of bacterial protector (Premax ® LLI, Rizobacter) (inoculant mixture). Seed inoculation was performed by immersing the seeds in the inoculant mixture for 60 min in 1.5 mL sterile tubes.

### 2.2 Plant growth evaluation on early vegetative stages

For all assays, tomato seeds (cv. UCO-14) were treated with *Streptomyces* sp. N2A (N2A) or water as control (C), both treatments with bacterial protector. For the germination assay, one hundred seeds per treatment were sown in four trays containing sand moistened with sterile distilled water and maintained in a chamber at 27°C with 12:12 h (light:dark) photoperiod. Seedlings emergence was evaluated at 7 days post sowing (dps). At 14 dps, normal and abnormal seedlings and non-germinated seeds were recorded according to ISTA rules (ISTA, 2022). The assay was repeated twice.

For plant vegetative growth and yield evaluations, after inoculation treated seeds were sown in seedling trays (55 cm^3^) containing peat substrate moistened with distilled water. Experiments were conducted under greenhouse conditions under a 14:10 h (light:dark) photoperiod, with a minimum temperature of 18°C and a maximum of 32°C. In the assay focused on vegetative stage, plants were transplanted to pots (300 cm^3^) containing a mixture of substrate (Kekkila®): perlite: dirt (1:1:1) at 30 dps, pots were sub-irrigated and arranged in a completely randomized design with 30 plants per treatment. The experiment was conducted for 50 dps, where the number of nodes per plant, stem middle perimeter and internode length for each treatment was recorded. At the end of the experiment, all plants were harvested by cutting the stem at the cotyledon node, oven-dried at 95°C for 96h, and weighed to determine dry biomass. The assay was performed in two independent experimental runs.

### 2.3 Fruit production and quality assessment

For yield evaluation, seedlings from N2A and C treatments were kept on trays until 30 dps and then transplanted to 10 L pots containing a mixture of substrate (Kekkila®): perlite: dirt (1:1:1) and maintained under the same greenhouse conditions described before. Pots were irrigated by a drip system three times per day and arranged in a completely randomized design with 30 plants per treatment. The assay was performed in two independent experimental runs.

Growth and development parameters including internode length (between the second and third node), stem perimeter (basal, middle, and apical), and number of flowers per inflorescence, were recorded for all plants at 40 days post-transplant (dpt). At the same time, in the onset of flowering and fruit set, leaf temperature and thickness were measured using a MultispeQ V2.0 device (Kuhlgert et al., 2016). During the following four weeks, the number of flowers and setting fruits were counted. At the end of the assay, fruits were periodically harvested at the red-ripe stage, defined as full red coloration of the fruit surface (Giovannoni, 2004). Each fruit was weighed using an analytical balance (Mettler Toledo ® ME 3002), and fruit shape was determined as the ratio height/diameter, measured with a manual caliper. Yield evaluation was defined at the end of the assay based on the total number of harvested fruits, individual and total weight per plant and treatment. Fruits showing visible disease symptoms were excluded from the final yield evaluation (Ozores-Hampton et al, 2015). After harvest, plants were cut down at the base and oven-dried to constant weight to evaluate biomass (dry weight) with a Kretz Delta2 scale.

### 2.4 Fruit quality analysis

Fruits used for quality analyses were harvested at the red-ripe stage, whereas fruits destined for postharvest shelf-life evaluation were harvested at the breaker stage (orange fruit). Fruit color was determined using a CR-400 chromameter (Minolta), with three measures taken at the equatorial zone of each fruit to calculate the a/b color index, where a and b correspond to absorbance values at 540 and 675 nm, respectively. Fruit firmness (scale from 0 to 100) was determined by two measurements at the equatorial zone using a Shore A durometer (Durofel DFT100) fitted with a 0.10 cm² tip. Fruits were then transversely sectioned to record the number of locules and pericarp thickness, measured with a manual caliper at five different points per fruit. Pericarps were homogenized to obtain fruit juice, which was used to measure pH, soluble solids content (°Brix) with a manual refractometer (SO-RH ®) and titratable acidity, calculated from the volume of 0.1 N NaOH required to titrate 10 g of juice diluted in 100 mL of distilled water to pH 8.1 (Rodríguez et al., 2005).

Postharvest shelf life (PSL) was evaluated by storing three fruits per plant at 25 °C on a dark chamber (Cambiaso et al., 2019). PSL was defined as the number of days from harvest until the appearance of deterioration symptoms, such as wrinkling and softening, for each treatment. Fruits were inspected every three days.

### 2.5 Microscopic analysis of ovary/fruit development around anthesis

An independent assay was conducted for flower and fruit microscopy and transcriptional analysis under the same conditions as those previously described. Thirty plants per treatment were arranged in a completely randomized design, and flower samples from each treatment (N2A and C) were collected at different times around anthesis: 2 days before anthesis (−2), at anthesis (0), and 3 and 7 days post-anthesis (dpa). Fifteen samples from each treatment and time point were taken and immediately fixed in a formaldehyde-acetic-ethanol (FAE) solution to preserve tissue structure. Subsequently, flowers samples were processed for histological analysis (ovary-fruit isolation) and cleared with methyl salicylate as described in Colono et al (2019). Briefly, fixed samples were dehydrated through an ethanol dilution series and transferred to pure methyl salicylate until tissues became translucent. Cleared tissues were mounted in methyl salicylate and examined under differential interference contrast (DIC) microscope (LEICA ® DM750 DIC). Images were acquired using the Leica Application Suite v 4.6.2 (Build:41.0) software. Pericarp thickness, number of cell layers and cell size were measured for each treatment and time point by observing four different parts of each fruit around the equatorial zone. Also, at each sampling time, tomato ovaries and fruits were visually examined using a hand lens (MIKOBA ®), and ovary size (length and width) was measured using IS-Caption Version 1.0 software ®. Growth rates were estimated for each variable by a regression analysis.

### 2.6 Gene expression analysis

In the same experiment described before, floral tissues from each treatment and sampling time (−2, 0, 3 y 7 dpa) from 10 independent plants were pooled in equal amounts (approximately 50 mg of isolated ovary-fruit), immediately frozen in liquid nitrogen, and used as one biological replicate for RNA extraction. Total RNA was extracted using TriPure® reagent (Roche). RNA quantification was measured with a NanoDrop Lite (ThermoScientific TM). Then, samples were treated with DNase I (ThermoFisher ®, Carlsbad, USA) and cDNA was synthesized using RevertAid reverse transcriptase (ThermoFisher ®, Carlsbad, USA) from 0.5 μg of RNA and dT18 oligonucleotide. After quality check of synthesized cDNA, it was used for the RT-PCR.

RT-PCR was made in a final volume of 10 μl containing 1μl of EVA Green (Biotium, San Francisco, USA), 10 pmol of each specific set of primer (Table S1), 3 mM MgCl_2_, cDNA dilution (1:10), and 1 U of Taq Polymerase (PBL, Buenos Aires, Argentina). Reaction was monitored with a Bio-Rad ® CFX Connect. Relative expression was calculated using the ΔΔCt method (Livak and Schmittgen, 2001). Relative gene expression was normalized against the Ubiquitin gene, used as housekeeping reference and the stability of this gene in each treatment and time was validated (Løvdal and Lillo, 2009). Gene expression was quantified from three biological replicates per evaluation time and analyzed with three technical replicates.

### Data analysis

Data was analyzed using R software (R Core Team, 2025) and Infostat (Di Rienzo, 2020). Variables expressed as proportions or counts derived from a fixed number of observations (binomial data) were analyzed using generalized linear mixed models assuming a binomial distribution. Continuous variables with a normality distribution were analyzed by analysis of variance (ANOVA) and a mean comparison test (Fisher’s LSD Test). Progression data was analyzed using a linear model and rate was calculated from the slope of the linear regression. Potential yield component trade-offs were evaluated using linear models relating individual fruit weight to fruit number. Yield components were analyzed by a Pearson’s correlation test to identify the variables contributing most to treatment differentiation. Statistical significance was set at *p* < 0.05.

## 3. RESULTS

### 3.1 Effects of bacterial inoculation on seed germination, seedling emergence and plant vegetative growth and development

Seeds inoculated with *Streptomyces* sp. N2A (N2A) showed a significantly enhanced seedling emergence compared to the control at 7 and 14 dps. No significant interaction was found between treatments and replicates. At 7 dps, N2A-treated seeds reached an emergence of 71.7 %, whereas control seeds exhibited an emergence of 55.1 %. Then, at 14 dps, N2A-inoculated seeds maintained a significantly higher emergence (83.2 %) than C (73.7 %). In addition, N2A treatment showed a significantly higher percentage of normal seedlings than control (Fig. 1a, b).

**Figure 1.**
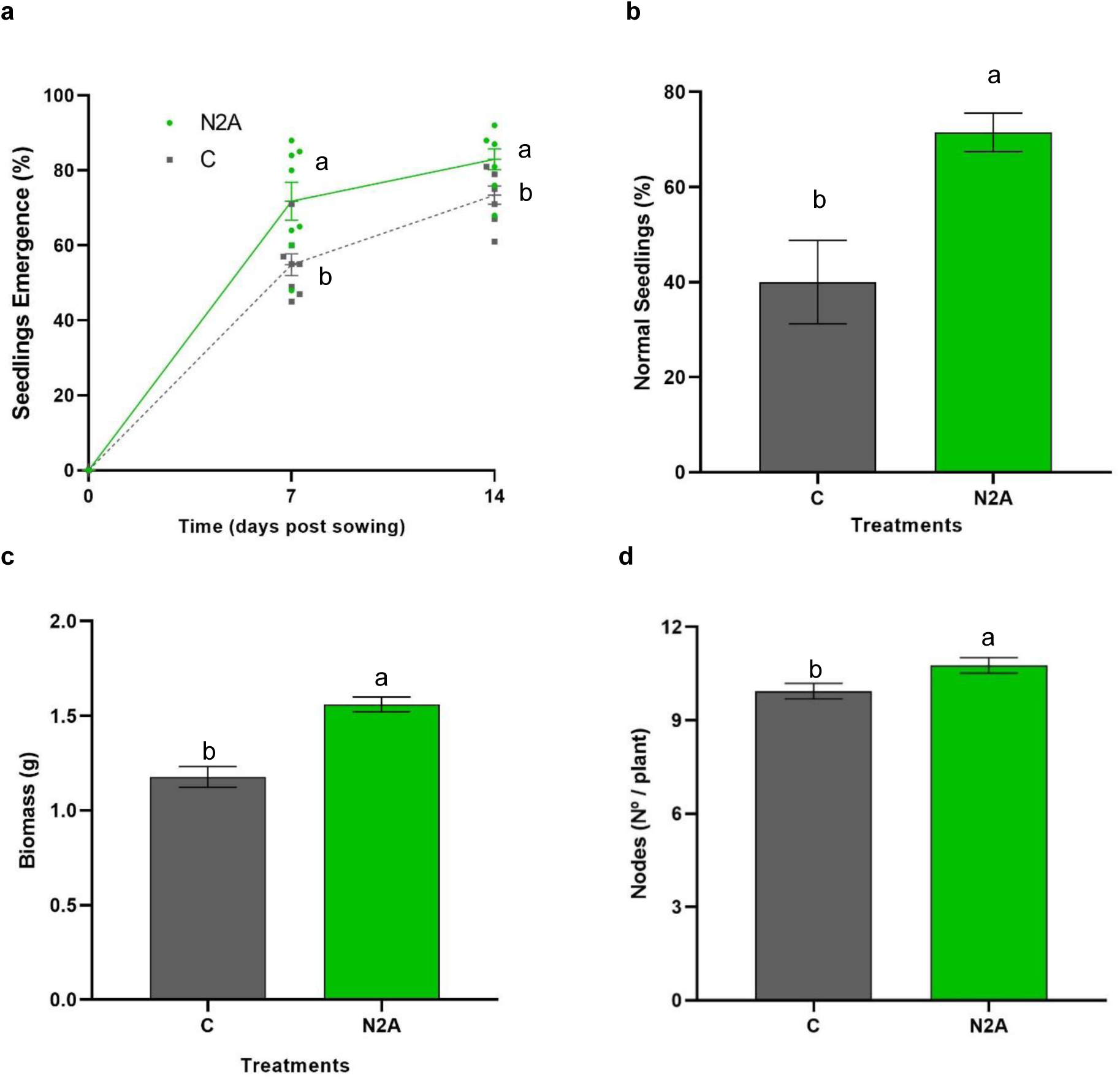
Effect of seed inoculation with *Streptomyces* sp. N2A (N2A in green) on tomato seedlings emergence and vegetative growth and development compared to control (C in grey). **(a)** Seedling emergence at 0, 7 and 14 days post sowing (dps), **(b)** normal seedling percentage at 14 dps, **(c)** biomass (dry weight) and **(d)** node (number per plant) at 50 dps. Different letters indicate significant differences between treatments according to a generalized linear mixed model and ANOVA followed by Fisher’s LSD Test (*p* < 0.05).

To further analyze the persistence of the *Streptomyces* sp. N2A effect, growth and development were evaluated during vegetative stage of tomato plants. At 50 dps, significant increments were found in the biomass accumulation (dry weight) and stem nodes number in N2A-treated plants, compared to C (Fig. 1c, d), indicating a growth and development promotion effect at early stages. No differences were found between treatments for the basal perimeter of the plant and number of flowers, but some differences in the medial stem perimeter were detected. Specifically, medial stem perimeter was higher on plants treated with N2A (2.89 cm) compared to control (1.42 cm), and consistently, N2A-treated plants presented a smaller internode length on this segment (between 3^rd^ and 4^th^ node) (2.32 cm) than control (3.20 cm) (Fig. S1).

### 3.2 Effect of *Streptomyces sp.* N2A over fruit development and yield

The fruit set dynamics over time showed no differences between treatments (Fig. S2). The cumulative fruit set rates were 1.91 and 2.12 average fruits per week for control and N2A respectively, while no treatment x time interaction was detected. There were no significant differences between regression slopes, indicating the absence of modifications in the rate of fruit production over time (Fig. S2). In addition, the analysis of leaf temperature differential (ΔT; leaf temperature minus air temperature) and leaf thickness values, at the fruit forming stage, showed no difference between treatments (Fig. S3).

Then, at harvest, yield was significantly increased in plants inoculated with *Streptomyces sp.* N2A in both years compared to C (*p* < 0.01), although there was a non-significant increment in fruit number (*p* = 0.120) (Fig. 2a, b). Thus, a significant increase in fruit weight was observed in plants treated with N2A, compared to C (*p* < 0.01) (Fig. 2b, c). No interaction in Treatment x experiment was found.

**Figure 2.**
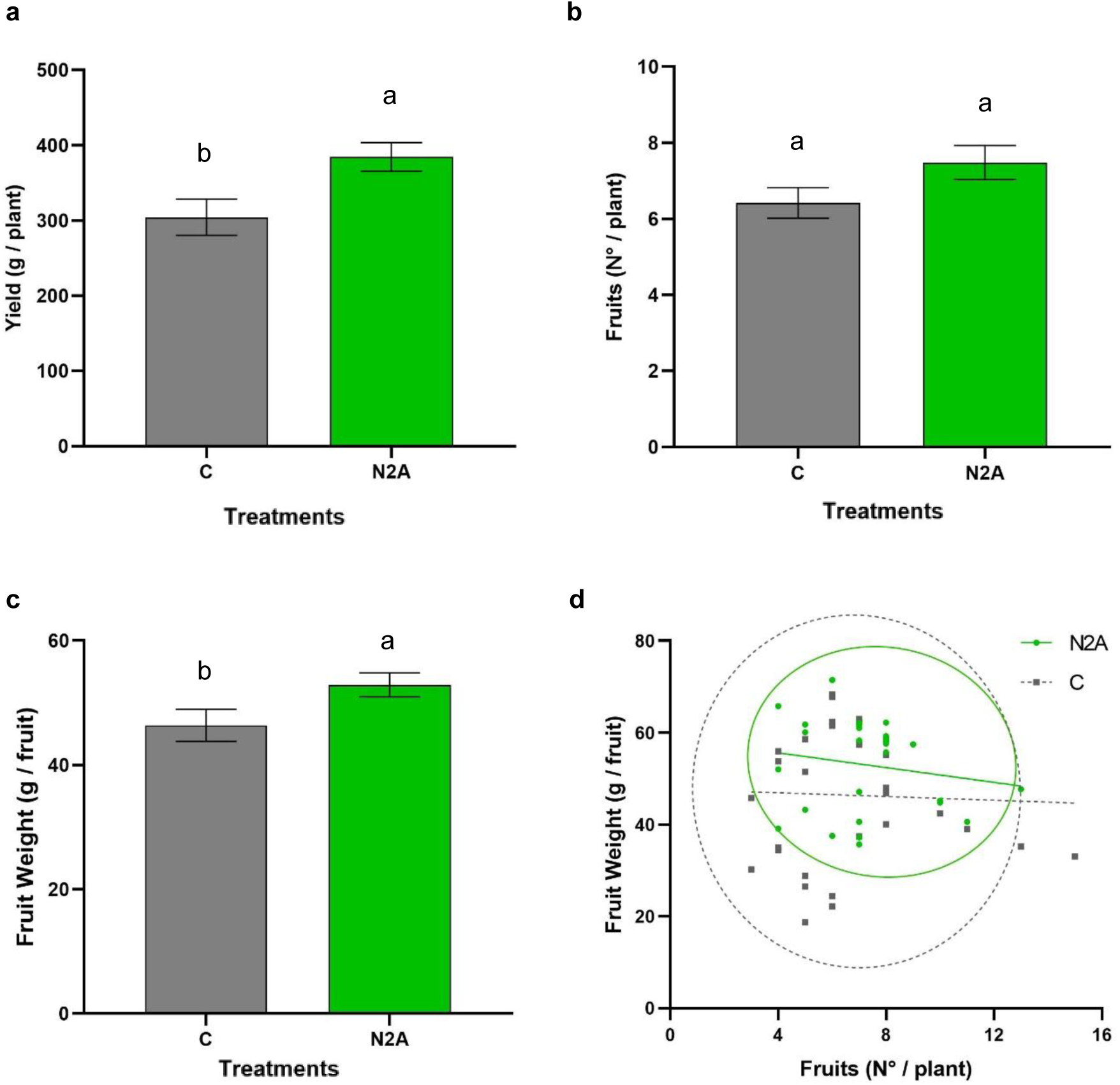
Yield components at harvest from both treatments, *Streptomyces sp.* N2A (N2A, green) and control (C, grey). **(a)** Mean yield expressed as grams per plant, **(b)** fruit number per plant, **(c)** individual fruit weight, and **(d)** trade-off relationship between fruit number and individual fruit weight. Points correspond to individual values. Different letters demonstrate significant differences between treatments according to an ANOVA followed by Fisheŕs LSD Test (*p* < 0.05). Trade-off was analyzed using linear models and slope test (*p* < 0.05). Ellipses represent the 95 % confidence region of each treatment according to Hotelling’s T^2^ approximation.

To evaluate potential trade-offs in yield components associated with plant resource partitioning and bacterial inoculation, the relationship between fruit number and individual fruit weight was analyzed. Individual fruit weight was not significantly related to fruit number and no significant interaction between fruit number and treatment was detected. Regression analyses showed slightly negative slopes between parameters, but with no significant differences between treatments (control slope = -0.20; N2A: slope = -0.95) (Fig. 2d). Moreover, based on these parameters, the N2A treatment significantly reduced the distance to the centroid, indicating lower phenotypic dispersion compared to control plants (*p* < 0.05) (Fig. 2d). Linear regression analyses further revealed that yield was significantly and positively associated with fruit number (r = 0.73, *p* < 0.01) and to individual fruit weight (R² = 0.64, *p* < 0.01).

### 3.3 Analysis of fruit morphology and quality

The analysis of fruit quality parameters, including pericarp firmness, pH, soluble solids content, and titratable acidity, showed no difference between fruits of N2A treated and control plants (Table 1). These results indicate that N2A treatment increased fruit weight and yield without significantly affecting the fruit quality parameters evaluated. In addition, no differences in color were found. Regarding fruit morphology, N2A treatment resulted in a significant increase in both pericarp thickness and fruit height compared with the control. Interestingly, a significantly different fruit shape was detected in response to the bacterial treatment due to an increase in fruit height but no differences in fruit width (Table 1).

**Table 1.** Mean values and standard error for tomato fruit morphology and quality parameters at harvest for both inoculation treatments: *Streptomyces sp.* N2A (N2A) and control (C).

|  | Firmness<br>(0 to 100) | pH | Soluble<br>solid<br>(°Brix) | Titrateable<br>acidity<br>(ml<br>NaOH) | Pericarp<br>thickness<br>(mm) | Size<br>(cm<sup>3</sup>) | Diameter<br>(mm) | Height<br>(mm) | Shape<br>index<br>(mm/mm) |
| --- | --- | --- | --- | --- | --- | --- | --- | --- | --- |
| N2A | 55.89 ±<br>1.77 a | 4.68 ±<br>0.03 a | 3.62 ±<br>0.08 a | 4.14 ±<br>0.14 a | 6.92 ±<br>0.16 a | 60.90 ±<br>3.26 a | 45.04 ±<br>1.07 a | 56.42 ±<br>1.36 a | 1.25 ± 0.03<br>a |
| C | 51.80 ±<br>2.09 a | 4.62 ±<br>0.02 a | 3.80 ±<br>0.10 a | 4.04 ±<br>0.12 a | 6.40 ±<br>0.18 b | 60.46 ±<br>3.70 a | 47.62 ±<br>1.10 a | 52.02 ±<br>1.40 b | 1.11 ± 0.03<br>b |
| Treatment | ns | ns | ns | ns | ** | ns | ns | * | ** |

Different letters indicate significant differences according to ANOVA LSD Fisher`s Test ( $p < 0.05$ ). In grey are highlighted the parameters with significant differences.
\* ( $p < 0.05$ ); \*\* ( $p < 0.01$ )
ns: non significant

Lastly, to further understand the correlation between evaluated parameters and pericarp thickness and to explain the basis of yield improvement, a Pearsońs correlation analysis was conducted with selected variables. One of the most significant and positive correlations was found between fruit weight and pericarp thickness (r = 0.55; *p* < 0.01) (Fig. 3a). Additionally, pericarp thickness was positively associated with fruit height (r = 0.67; *p* < 0.01), diameter (r = 0.25; *p* = 0.143) and size (r = 0.49; *p* < 0.01) (Fig. 3b-d). Nevertheless, a moderate negative correlation was found between pericarp thickness and fruit number (r = −0.47; *p* < 0.05) (Fig. 3e). Finally, pericarp thickness was not directly associated with yield (r = 0.02; *p* = 0.8869) (Fig. 3f).

**Figure 3.**
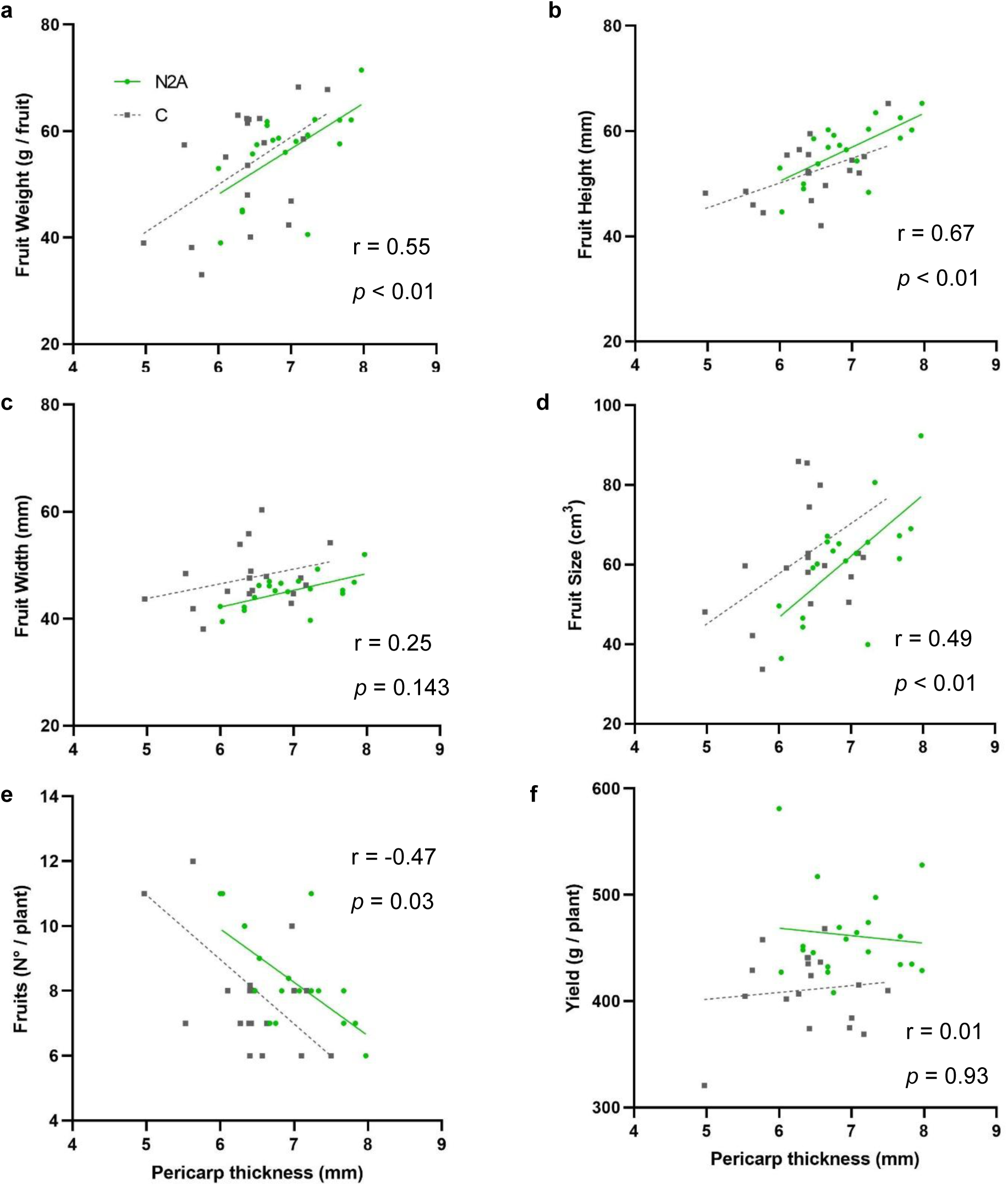
Associations between pericarp thickness and (**a**) fruit weight, (**b**) fruit height, (**c**) fruit diameter, (**d**) fruit size, (**e**) fruit number, and (**f**) yield; for both treatments: *Streptomyces* sp. N2A (N2A, green) and control (C, grey). Lines indicate linear regression of both treatments separately. Pearson correlation coefficients (r) and *p*-values were calculated from all observations pooled across treatments.

### 3.4 Microscopic changes in ovary to fruit transition induced by *Streptomyces sp.* N2A

The early effects of bacterial inoculation on ovary and fruit morphology (size and pericarp thickness) were further evaluated through an additional experiment. Based on microscope assessments of overall morphology and a regression analysis, no differences were found in the growth rate of ovary/fruit width, height and size between treatments (Fig. 4; Fig. 5a-c; Table S2). However, fruits from the N2A-treated plants were significantly higher from 0 dpa onwards, on each time point (Fig. 4; Fig. 5a; Table S2).

**Figure 4.**
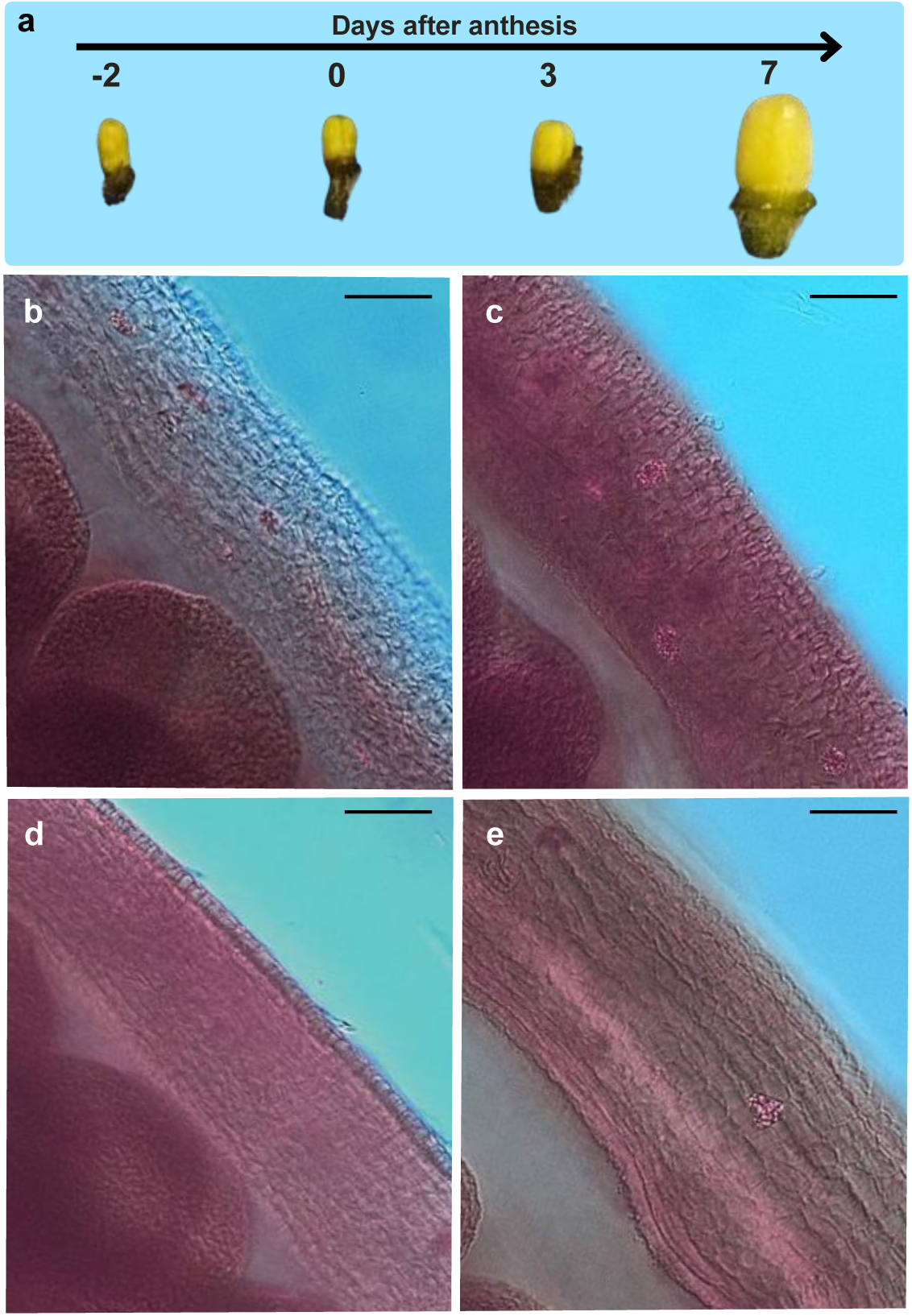
Development of ovary/fruit traits and pericarp cellular parameters in both treatments: *Streptomyces sp.* N2A and water as control (C) before (−2), 7 days post anthesis. **(a)** Overall progression of ovary/fruit growth; **(b)** C pericarp at −2 and **(c)** 7 dpa; **(d)** N2A pericarp at −2 and **(e)** 7 dpa. Bar represents 50 μm under microscope visualization at 50x.

**Figure 5.**
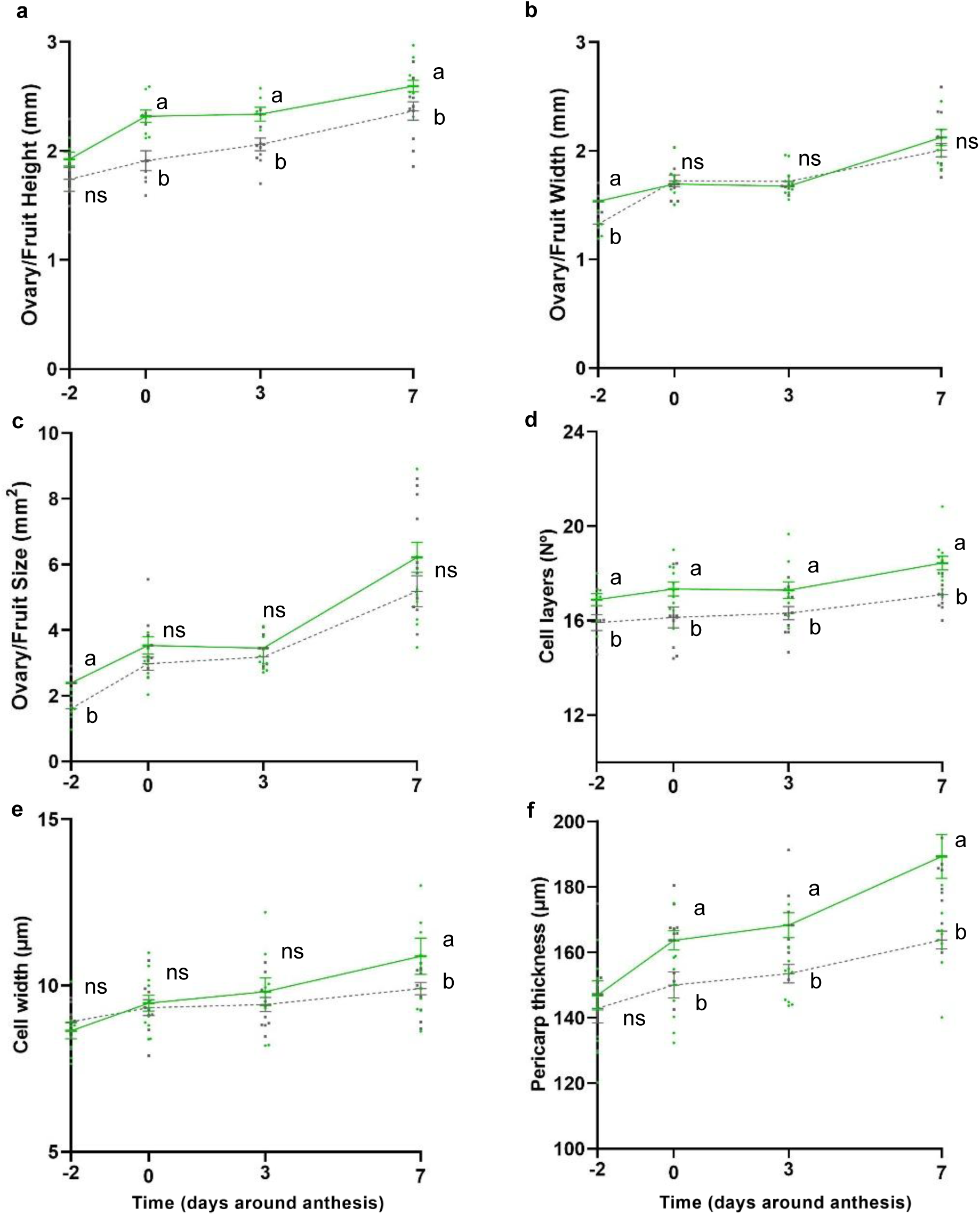
Development of ovary/fruit traits in both treatments: *Streptomyces* sp. N2A (green) and water (C, grey) before (−2), at anthesis (0) and post anthesis (3 and 7 days post anthesis, dpa). (**a**) ovary-fruit height (mm), (**b**) ovary-fruit width (mm), (**c**) ovary-fruit size (mm³), (**d**) cell layers (N°), (**e**) cell width (µm) and (**f**) pericarp thickness (µm). Points represent individual observations; lines connect treatment means and error bars indicate mean ± standard error. Differences in developmental slopes between treatments were evaluated by linear regression, while treatment differences at each time point were assessed using ANOVA followed by Fisher’s LSD test (p < 0.05). ns: non significative.

Regression analysis showed no differences in the rate of cell layers development between treatments, and a non-significant increase in the rate of cell expansion (cell width) was observed over time (*p*=0.117) (Fig. 4; Fig. 5d-e; Table S2). Consistently, fruits from N2A-treated plants showed a significantly higher number of cell layers than C since −2 dpa (Fig. 4; Fig. 5d; Table S2). Thus, pericarp thickness showed the most pronounced differences, being significantly thicker in fruits from N2A-treated plants since 0 dpa and displaying a significantly greater growth rate compared to C, indicating profound differences in fruit growth and development from anthesis onwards (*p* < 0.05) (Fig. 4; Fig. 5f; Table S2). Among the overall fruit morphological traits, fruit height and pericarp thickness were the only two parameters showing significant differences between treatments from anthesis onwards (Fig. 4; Fig. 5a, f; Table S2).

### 3.5 Gene expression analysis in ovary to fruit transition

The qRT-PCR analysis was conducted during the ovary to fruit transition to analyze different hormonal signaling pathways and cell cycle-related genes potentially involved in the observed increase in fruit weight and pericarp thickness. Genes related to auxin signaling and biosynthesis such as *IAA17* and *TOFZY2*, respectively, were found to be repressed in ovaries/fruit from N2A-treated plants at −2 dpa and induced at 7 dpa compared to the control (Fig. 6; Fig. S4). Consistently, the *ARP* gene, encoding an auxin-repressed protein, was found to be induced at −2 dpa but markedly repressed at 7 dpa compared to C. Moreover, genes associated with cytokinin signaling, such as *CKX2* and *ZEATIN* were found to be differentially expressed, being strongly repressed and induced, respectively, at −2 and 0 dpa in N2A-treated plants compared to C and were later induced at 7 dpa. The *GAMYB2* gene, associated with the gibberellin pathway, was less affected by bacterial treatment, showing a slight induction at −2 dpa and repression at 0 dpa (Fig. 6; Fig. S4).

**Figure 6.**
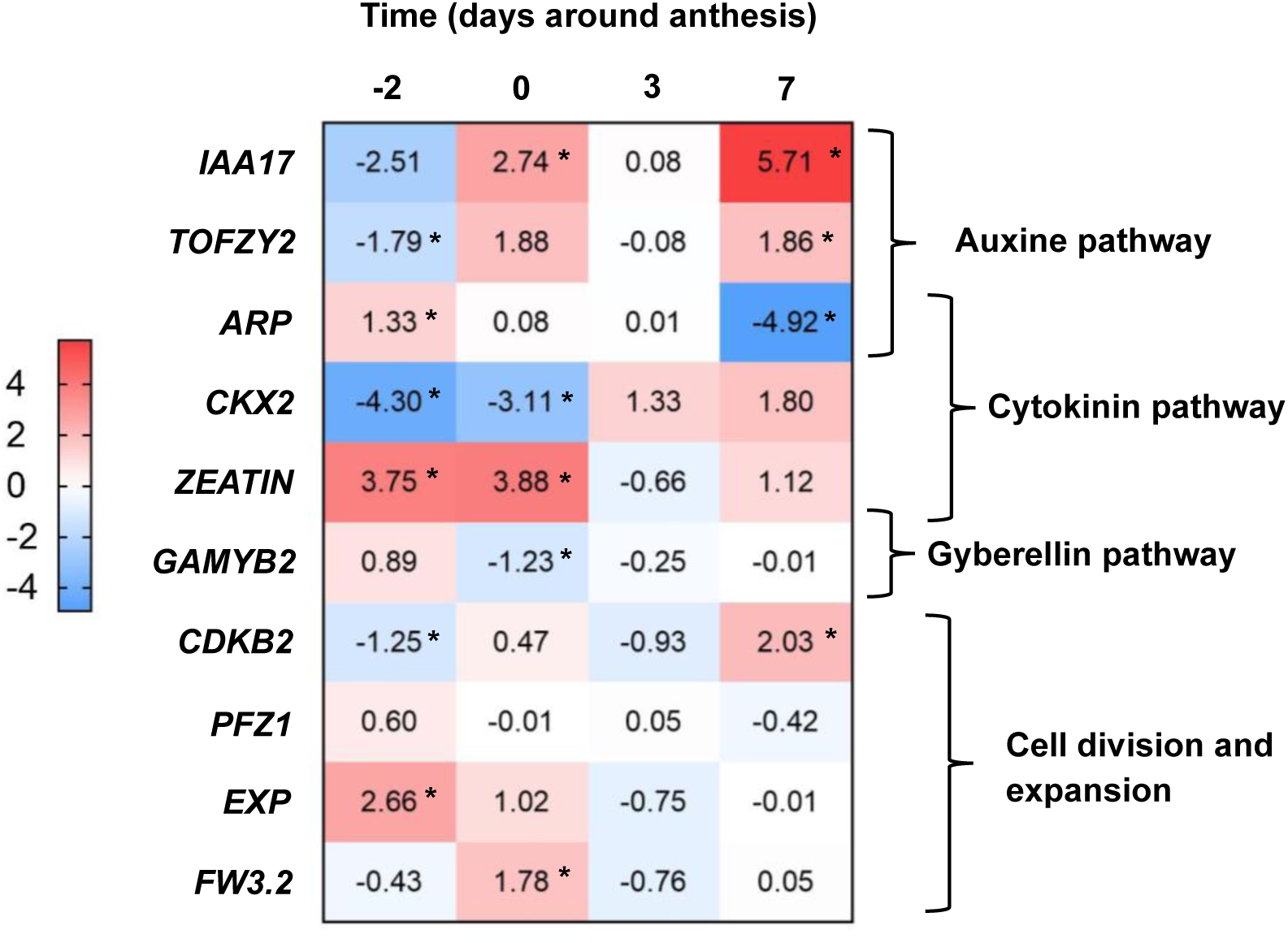
Heat map showing the relative expression of induced (red) and repressed (blue) genes related to hormonal signaling and cell division and expansion during early ovary to fruit transition at 2 days before anthesis (−2), anthesis (0) and 3 and 7 days post anthesis, determined by qRT-PCR. Expression values represent the Log fold change of genes in tissue from *Streptomyces* sp. N2A treatment, relative to control (water treatment) at each time point and were normalized using Ubiquitin as the internal reference gene. Significant differences between treatments at each time point were determined using ΔCt values by ANOVA followed by Fisher’s LSD Test. (*) indicates significant differences at *p* < 0.05.

The expression analysis of genes encoding proteins associated with cell division and expansion, such as the *PFZ1* and *FW3.2* genes, resulted in minor differences between treatments, throughout the evaluated time points. The cyclin-dependent kinase gene *CDKB2* showed a slight repression at −2 dpa but a strong induction at 7 dpa. Meanwhile, the *EXP* gene encoding an expansine protein associated with cell wall loosening, was strongly induced in N2A-treated plants at −2 and 0 dpa (Fig. 6; Fig. S4).

## 4. DISCUSSION

In recent years, a growing number of studies have focused on the effects of different PGPR on plant performance, particularly highlighting their roles in enhancing plant growth, diseases biocontrol and abiotic stress tolerance. The results showed in this work reinforce previous findings around the capacity of *Streptomyces spp.* and other PGPRs to promote seed germination, seedling emergence and growth and development of tomato plants at early vegetative stage, including increased biomass production and nodes number (Sayyed et al, 2024; Rehan et al, 2023; Kawicha et al, 2024). These effects are commonly associated with auxin biosynthesis and transport induction by different rhizobacteria, which may directly stimulate plant cell elongation and development (Rehan et al, 2023). Accordingly, *Streptomyces* sp. N2A was previously characterized as an Indole Acetic Acid (IAA) producer strain, among other plant growth-promoting traits, showing also a positive impact in soybean seed germination, seedling emergence and biomass (Villafañe et al., 2024). Similarly, in the present study, N2A-treatment promoted vegetative growth in tomato, indicating that its plant growth promoting *Streptomyces* (PGPS) effects are consistent across both crops. However, the analysis of the effects of N2A inoculation on the flowering dynamics in tomato plants at the reproductive stage showed no differences at the flowering-rate between treatments; in contrast with the higher reproductive nodes observed in soybean (Villafañe et al, 2024); indicating that there are also some differences in its behavior among crops.

The impact of PGPR inoculation on tomato yield has been previously evaluated in other works, mainly reporting increases in fruit number and individual weight (Katsenios et al, 2021; Gashash et al, 2022; Ogugua et al, 2025). Consistently, *Streptomyces* sp. N2A inoculation was previously shown to increase reproductive node number, pod number and consequently yield in soybean crop under field conditions, without affecting seed unitary weight (Villafañe et al. 2024). In the tomato cultivar evaluated here, yield improvement was associated with increased fruit weight rather than fruit number; suggesting that the yield component affected may vary among crops, genotypes and growth conditions. These results partially align with those of Khomampai et al. (2024), who observed increases in individual fruit weight and size in tomato plants treated with actinomycetes. However, they reported a negative correlation between fruit number and fruit weight, which was not detected in our study. As expected, yield was positively associated with both fruit number and individual weight. Additionally, pericarp thickness, another important fruit morphological trait, is often moderate to strongly associated with fruit size and weight. Susic et al. (2000) reported consistent correlations among these traits and found that they increased significantly following bacterial treatment. Besides, in our work, no differences were found in several parameters associated with fruit quality like pH, soluble solids or titratable acids, indicating that the increment in fruit weight does not compromise potential nutritional value of the tomato fruits.

In this work, it was found that *Streptomyces* sp. N2A inoculation on tomato seeds modified fruit pericarp thickness and its association with yield components. Notably, the differences in the pericarp observed at harvest in response to N2A inoculation were already detectable during the ovary/fruit growth and development around anthesis, highlighting a critical gap in the understanding of the effects of this PGPS at this developmental stage. Thus, significant differences were found in ovary height and pericarp thickness, consistent with those observed at harvest. Given the absence of significant differences in fruit number, the present study focused on fruit morphology traits to investigate the mechanisms underlying the N2A mediated increase in yield. This approach was supported by evidence that tomato fruit weight and size are largely determined during early fruit development and are ultimately associated with the final yield (Frary et al, 2000; Giovannoni 2004; Beauchet et al, 2024).

Tomato fruit morphology results from the coordinated action of genetic factors and hormone-mediated regulatory pathways. Auxins, gibberellins, ethylene, and brassinosteroids (BRs) contribute to the regulation of fruit size and shape through interconnected developmental mechanisms (Lazaro et al. 2018; Rafiq at al. 2025). On the other hand, it is noteworthy that no previous work has specifically examined the modulation of phytohormonal pathways by PGPSs during the ovary-to-fruit transition in tomato. Therefore, in the present study, to further understand the mechanisms underlying the effects of *Streptomyces* sp. N2A inoculation on fruit establishment, particularly in pericarp thickness and consequently weight, a gene expression analysis of key hormonal and cell cycle-related pathways was conducted during early fruit development, providing novel evidence of PGPS effect at this critical stage. Current knowledge of genetic and molecular mechanisms controlling the shape of harvestable organs has been derived largely from studies in rice and tomato. In rice, changes in grain size are frequently accompanied by alterations in shape and involve regulators from multiple pathways, including G-protein signaling, ubiquitin-mediated protein degradation, phytohormone signaling, particularly BRs, auxins, and cytokinins, and transcriptional control. By contrast, the proteins identified in tomato appear to represent a narrower functional range and are mainly associated with cytoskeletal organization and cell-division patterns (Lazzaro et al., 2018).

The phytohormones pathways evaluated in the present study have a well-documented role in fruit development; since reduced cytokinin levels are significantly correlated with smaller fruit size, lower fruit weight and thinner pericarp (Gan et al, 2022). In addition, gibberellins have a critical impact on cell size, cell layers number and endoreduplication (Renaudin et al, 2023); and auxins signaling, transport and endogenous levels directly affect cell size, pericarp thickness and fruit set by regulating cell division and expansion (Sundaresan et al, 2016; Liu et al, 2018; Zhang et al, 2021). Therefore, to further understand which biochemical pathways were induced or modified by the bacterial inoculation, expression of several genes was evaluated during this period.

Within the auxin pathway, *IAA17* gene, which encodes a transcriptional repressor induced by elevated auxin levels in the pericarp and act as a negative regulator of auxin signaling (Mastoraki et al, 2025), was upregulated in tissue from N2A treatment around anthesis and at 7 dpa. Its expression pattern was positively associated with that of *TOFZY2*, which encodes a key enzyme involved in auxin biosynthesis, particularly in ovules after pollination (Zhang et al, 2016). Consistent with the auxin dynamics described above, the *ARP* gene, encoding an auxin-repressed protein expressed in reproductive tissues, showed reduced expression as the auxin levels increase, as previously reported (Cheng et al, 2025; Gan et al, 2022; Ma et al, 2025). These changes in auxin biosynthesis and signaling may also be relevant to the morphological differences observed during ovary and early fruit development. Auxin has been proposed to influence microtubule organization through *SUN/IQ67-DOMAIN* (IQD)-mediated pathways, thereby linking hormonal signaling with cytoskeletal dynamics (Lazzaro et al., 2018). Through this mechanism, auxin signaling could alter cytoskeletal dynamics and thereby influence oriented cell division and cell expansion, two processes that contribute to organ shape. Although IQD proteins and microtubule dynamics were not directly evaluated here, this pathway provides a plausible link between the N2A-associated changes in auxin regulation and the differences detected in ovary and fruit morphology.

Meanwhile, cytokinin play important roles in cell division and fruit set and may also modulate *ARP,* as this gene has been reported to be upregulated under cytokinin-deficient cultivars (Gan et al, 2022). In this pathway, *CKX2*, codifying a cytokinin degradation enzyme, is strongly repressed before and during anthesis (−2 and 0 dpa) compared to C, indicating that cytokinins are not being degraded at the same rate into irreversible forms in tissue from N2A treatment and at these stages (Baranov and Timerbaev, 2024). At these two evaluated time points, *Zeatin*, encoding a zeatin O-glucosyltransferase-like protein, was overexpressed in tissue from N2A-treated plants, which is associated with increased Zeatin levels, due to its ability to store cytokinin into reversible inactivated forms and its resistance to cytokinin oxidase activity (*CKX2*) (Kieber and Schaller, 2014; Chen et al, 2021). After anthesis, when auxin-pathway take the leading role in promoting fruit growth, *Zeatin* relative expression decrease and *CKX2* is induced in N2A-treated tissue compared to C, likely reflecting a negative feedback mechanism in response to elevated cytokinin levels in the growing fruit (Schmülling et al, 2003: Matsuo et al, 2012).

Regarding the gibberellin-pathway, *Gamyb2*, a MYB transcription factor, was found to be slightly induced before anthesis and repressed at anthesis in N2A-treated tissue. These small and dynamic changes could be explained by a limited bacterial effect on gibberellin metabolism. The repression at 0 dpa would be related to the induced activation of auxin biosynthesis, as *Gamyb2* acts as a negative regulator of ovary growth at this developmental stage (Wang and Maagd, 2025).

In addition, regarding cell division, the *CDKB2* gene was evaluated. This gene encodes a cyclin dependent kinase involved in the regulation of cell division (Czerednik et al, 2012). Accordingly, the overexpression of *CDKB2* resulted in smaller fruits due to a disruption in cell division and expansion. In ovaries from N2A-treated plants *CDKB2* was slightly repressed at 3 dpa and induced at 7 dpa, compared to C. The later overexpression could be associated with a maintenance of mitotic activity in the pericarp compared to C, due to higher level of auxins (de Jong et al 2009; Czerednik et al, 2012).

No consistency between treatments were detected in the expression of *PFZ1*, which encodes a zinc finger-domain protein, or *FW3.2/SlKLUH,* a major determinant of fruit weight and size. Nevertheless, *FW3.2*, showed a moderate, transient induction in ovaries from N2A-treated plants at 0 dpa, in agreement with the induction of auxin and cytokinin related pathways at this stage (de Jong et al, 2009; Rodríguez et al, 2011). However, natural variation at *FW3.2/SlKLUH* does not appear to affect ovary size at anthesis, as its effect on fruit weight becomes evident only several weeks after anthesis. This later effect has been attributed mainly to an extended period of cell division, leading to increased cell numbers and larger pericarp and septum tissues (Chakrabarti et al., 2013). Therefore, the transient induction observed at 0 dpa may represent an early molecular response to N2A treatment rather than an immediate effect on ovary morphology.

Lastly, *EXP*2, an expansin-encoding gene involved in cell enlargement and pectin modifications, was induced at −2 and 0 dpa, suggesting an early activation of cell wall loosening (Sundaresan et al, 2016). This effect is consistent with the increased pericarp thickness and cell size observed in the N2A-treated plants. The early induction of *EXP2* could also be associated with the induction of the auxin biosynthesis and signaling, as auxin is a well-known inducer of *EXP* expression during fruit growth (Catalá et al, 2000). Altogether, the gene-expression patterns observed during the ovary-to-fruit transition suggest a coordinated modification of the hormonal crosstalk induced by *Streptomyces* sp. N2A seed-inoculation over fruit development. However, further studies will need to be conducted to determine more precisely the exact mechanisms involved.

Overall, seed inoculation with *Streptomyces* sp. N2A improved several agronomic traits throughout the tomato crop cycle (Fig. 7). During the early vegetative developmental stage, this PGPS enhanced seed germination, seedlings emergence, and vegetative growth, likely through the direct and indirect plant growth-promoting traits previously characterized in this strain (Villafañe et al., 2024). Then, at reproductive stage, an increased auxin biosynthesis and signaling in the ovary/fruit, together with an induced cytokinin metabolism prior to anthesis, would contribute to an increased number of cells, cell-layers and enhanced cell expansion in the pericarp in the ovary-to fruit transition. These hormonal and morphological changes were observed from anthesis onwards and resulted in the thicker pericarp and consequently increased fruit weight observed at harvest, without affecting fruit quality (Fig. 7). Collectively, these findings indicate that the beneficial effects of N2A seed inoculation persist across successive developmental stages and ultimately contribute to increasing crop productivity.

**Figure 7.**
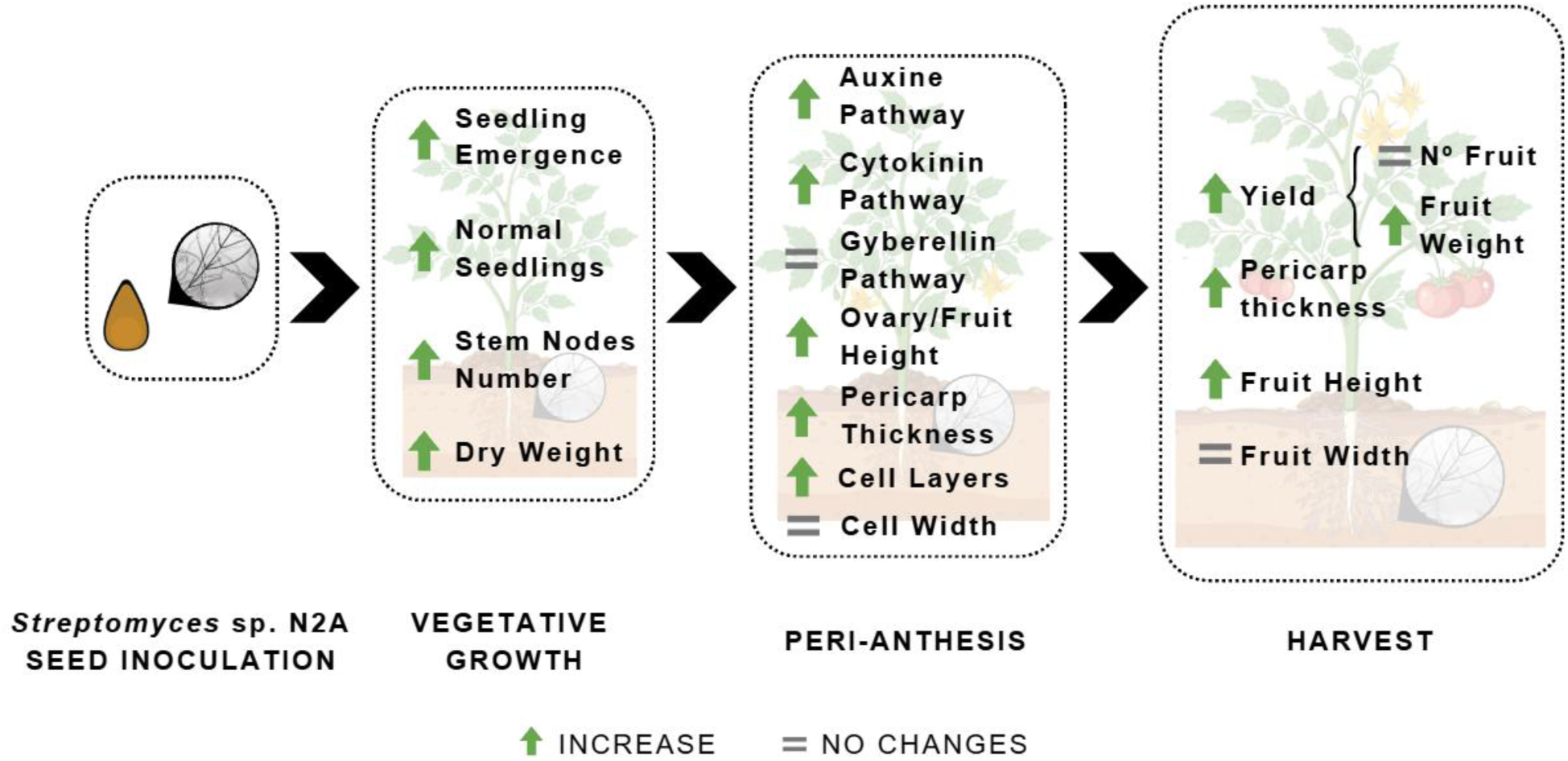
Summary diagram of the effects of *Streptomyces* sp. N2A (N2A) inoculation, compared with the control (C), on tomato vegetative growth, ovary-to-fruit development and yield components. Green upward arrows indicate a significant increase in response to N2A inoculation, whereas the equal sign indicates no significant differences between treatments.

## 5. CONCLUSIONS

This study provides the first evidence that a PGPS, particularly the strain *Streptomyces* sp. N2A, can increase fruit weight and consequently yield, by promoting pericarp thickness without affecting the fruit quality parameters evaluated. These effects may result from the modulation of multiple hormonal and cell wall-modifying enzyme pathways during the ovary-to-fruit transition in tomato. Besides, N2A-seed inoculation improves several agronomic traits from the earliest stages of tomato plant growth and development, promoting normal seedlings emergence and several biometric parameters at vegetative stage. Therefore, this bio-stimulant strain, isolated from soybean plant’s rhizosphere, would also be used in sustainable tomato production and establishes a novel reference framework for future research in PGPS-mediated mechanisms in fruit development.

## Acknowledgments

We thank to INTA La Consulta for providing tomato cv. UCO-14 seeds, to Dr. Alvaro Quijano for providing the PhotosynQ MultispeQ V2.0, and to Ing. Daniel Faura and David Balavan (IICAR) for providing technical assistance in greenhouse assays. We thank the Ministry of Environment and Climate Change of the province of Santa Fe, Argentina, for granting the corresponding permit for access and/or use of provincial natural resources, according to Prov. Res. No. 125/19.

## Funding

This study was supported by grants from Agencia Nacional de Promoción Científica y Tecnológica (ANPCyT) PICT-StartUp 2020-00038, Universidad Nacional de Rosario, and Consejo Nacional de Investigaciones Científicas y Técnicas (CONICET) PIP 11220220100491CO. R. Maldonado was awarded a scholarship by CONICET. M.A. Chiesa, E. Rodríguez and G. Rodríguez are CONICET Career Researchers.

## Author contribution

R.M., M.A.C., E.R. and G.R designed the research. R.M. and O.I. performed the assays. R.M. and M.A.C analyzed the data and wrote the first version of the manuscript. E.R. and G.R reviewed the manuscript. Funding was provided by grants to E.R. and M.A.C. All authors contributed to manuscript revision, read and approved the submitted version.

## Declaration of Competing Interest

The authors declare that they have no conflict of interest.

## Notes

### Competing Interest Statement

The authors have declared no competing interest.

## REFERENCES

1. Ali, M., Kamran, M., Abbasi, G. H., Saleem, M. H., Ahmad, S., Parveen, A., … & Fahad, S. (2021). Melatonin-induced salinity tolerance by ameliorating osmotic and oxidative stress in the seedlings of two tomato (Solanum lycopersicum L.) cultivars. Journal of Plant Growth Regulation, 40(5), 2236–2248.

2. Baranov, D., & Timerbaev, V. (2024). Recent advances in studying the regulation of fruit ripening in tomato using genetic engineering approaches. International Journal of Molecular Sciences, 25(2), 760.

3. Beauchet, A., Bollier, N., Grison, M., Rofidal, V., Gévaudant, F., Bayer, E., … & Chevalier, C. (2024). The cell number regulator FW2. 2 protein regulates cell-to-cell communication in tomato by modulating callose deposition at plasmodesmata. Plant Physiology, 196(2), 883–901.

4. Bhandari, P., Kim, J., & Lee, T. G. (2023). Genetic architecture of fresh-market tomato yield. BMC Plant Biology, 23(1), 18.

5. Cambiaso V, Gimenez MD, Vazquez VD, Pereira da Costa JH, Picardi LA, Pratta GR, Rodríguez GR (2019) Selected genome regions for fruit weight and shelf life in tomato RILs discernible by markers based on genomic sequence information. Breeding Science 69:447–454.

6. Catalá, C., Rose, J. K., & Bennett, A. B. (2000). Auxin-regulated genes encoding cell wall-modifying proteins are expressed during early tomato fruit growth. Plant physiology, 122(2), 527–534.

7. Chen, L., Zhao, J., Song, J., & Jameson, P. E. (2021). Cytokinin glucosyl transferases, key regulators of cytokinin homeostasis, have potential value for wheat improvement. Plant biotechnology journal, 19(5), 878–896.

8. Cheng, Y. J., Zhang, Y., Yu, J., Meng, X. R., Chang, Z. Y., Jiang, F. L., … & Zhang, T. Q. (2025). Chromatin accessibility analysis reveals functional cis regulatory regions related to fruit development and domestication in tomato. The Plant Journal, 123(3), e70416.

9. Colono, C., Ortiz, J. P. A., Permingeat, H. R., Souza Canada, E. D., Siena, L. A., Spoto, N., … & Pessino, S. C. (2019). A plant-specific TGS1 homolog influences gametophyte development in sexual tetraploid Paspalum notatum ovules. Frontiers in Plant Science, 10, 1566.

10. Cong, B., Liu, J., & Tanksley, S. D. (2002). Natural alleles at a tomato fruit size quantitative trait locus differ by heterochronic regulatory mutations. Proceedings of the National Academy of Sciences, 99(21), 13606–13611.

11. Czerednik, A., Busscher, M., Bielen, B. A., Wolters-Arts, M., de Maagd, R. A., & Angenent, G. C. (2012). Regulation of tomato fruit pericarp development by an interplay between CDKB and CDKA1 cell cycle genes. Journal of Experimental Botany, 63(7), 2605–2617.

12. da Silva, E. M., Silva, G. F. F. E., Bidoia, D. B., da Silva Azevedo, M., de Jesus, F. A., Pino, L. E., … & Nogueira, F. T. S. (2017). micro-RNA 159 targeted Sl GAMYB transcription factors are required for fruit set in tomato. The Plant Journal, 92(1), 95–109.

13. de Andrade, L. A., Santos, C. H. B., Frezarin, E. T., Sales, L. R., & Rigobelo, E. C. (2023). Plant growth-promoting rhizobacteria for sustainable agricultural production. Microorganisms, 11(4), 1088.

14. De Jong, M., Mariani, C., & Vriezen, W. H. (2009). The role of auxin and gibberellin in tomato fruit set. Journal of experimental botany, 60(5), 1523–1532.

15. De Jong, M., Wolters Arts, M., Feron, R., Mariani, C., & Vriezen, W. H. (2009). The Solanum lycopersicum auxin response factor 7 (SlARF7) regulates auxin signaling during tomato fruit set and development. The Plant Journal, 57(1), 160–170.

16. Di Rienzo, J., Balzarini, M., Gonzalez, L., Casanoves, F., Tablada, M., & Walter Robledo, C. (2010). Infostat: software para análisis estadístico.

17. Drobek, M., Frąc, M., & Cybulska, J. (2019). Plant biostimulants: Importance of the quality and yield of horticultural crops and the improvement of plant tolerance to abiotic stress—A review. Agronomy, 9(6), 335.

18. Food and Agriculture Organization of the United Nations (FAO). 2025. Agricultural Production Statistics 2010–2025. December 2025 update. FAOSTAT. Rome, Italy. Available at: https://www.fao.org/statistics/events/events-detail/agricultural-production-statistics-2010-2025.-december-2025-update/en

19. Frary, A., Nesbitt, T. C., Frary, A., Grandillo, S., Knaap, E. V. D., Cong, B., … & Tanksley, S. D. (2000). fw2. 2: a quantitative trait locus key to the evolution of tomato fruit size. Science, 289(5476), 85–88.

20. Gan, L., Song, M., Wang, X., Yang, N., Li, H., Liu, X., & Li, Y. (2022). Cytokinins are involved in regulation of tomato pericarp thickness and fruit size. Horticulture Research, 9, uhab041.

21. Gashash, E. A., Osman, N. A., Alsahli, A. A., Hewait, H. M., Ashmawi, A. E., Alshallash, K. S., … & Ibrahim, M. F. (2022). Effects of plant-growth-promoting rhizobacteria (PGPR) and cyanobacteria on botanical characteristics of tomato (Solanum lycopersicon L.) plants. Plants, 11(20), 2732.

22. Giovannoni, J (2004). Genetic Regulation of Fruit Development and Ripening, The Plant Cell, 16, (1)170–180, 10.1105/tpc.019158

23. Higashide, T. (2009). Prediction of tomato yield on the basis of solar radiation before anthesis under warm greenhouse conditions. HortScience, 44(7), 1874–1878.

24. International Seed Testing Association (ISTA). 2022. International Rules for Seed Testing. Effective from 1 January 2022. International Seed Testing Association, Bassersdorf, Switzerland. 10.15258/istarules.2022.I

25. Katsenios, N., Andreou, V., Sparangis, P., Djordjevic, N., Giannoglou, M., Chanioti, S., … & Efthimiadou, A. (2021). Evaluation of plant growth promoting bacteria strains on growth, yield and quality of industrial tomato. Microorganisms, 9(10), 2099.

26. Kawicha, P., Nitayaros, J., Sangdee, K., Thanyasiriwat, T., Somtrakoon, K., & Sangdee, A. (2024). Genomic insights into Streptomyces hygroscopicus subsp. hygroscopicus SRF1: a potential biocontrol agent against fusarium wilt with plant growth-promoting abilities in tomatoes. Biocontrol Science and Technology, 34(5), 389–410.

27. Khomampai, J., Jeeatid, N., Kaeomuangmoon, T., Pathom-Aree, W., Rangseekaew, P., Yosen, T., … & Chromkaew, Y. (2024). Endophytic actinomycetes promote growth and fruits quality of tomato (Solanum lycopersicum): an approach for sustainable tomato production. PeerJ, 12, e17725.

28. Kieber, J. J., & Schaller, G. E. (2014). Cytokinins. The Arabidopsis Book/American Society of Plant Biologists, 12, e0168.

29. Kim, D., Kim, J., Lee, Y., Balaraju, K., Hwang, Y. J., Lee, M. H., … & Jeon, Y. (2024). Evaluation of Streptomyces sporoverrucosus B-1662 for biological control of red pepper anthracnose and apple bitter rot diseases in Korea. Frontiers in Microbiology, 15, 1429646.

30. Kuhlgert, S., Austic, G., Zegarac, R., Osei-Bonsu, I., Hoh, D., Chilvers, M. I., … & Kramer, D. M. (2016). MultispeQ Beta: a tool for large-scale plant phenotyping connected to the open PhotosynQ network. Royal Society open science, 3(10), 160592.

31. Lankinen, Å., Witzell, J., Aleklett, K., Furenhed, S., Karlsson Green, K., Latz, M., … & Grenville-Briggs, L. (2024). Challenges and opportunities for increasing the use of low-risk plant protection products in sustainable production. A review. Agronomy for Sustainable Development, 44(2), 21.

32. Lazzaro M.D., Wu S., Snouffer A., Wang Y. & van der Knaap E. (2018). Plant Organ Shapes Are Regulated by Protein Interactions and Associations With Microtubules. Front. Plant Sci. 9:1766. doi: 10.3389/fpls.2018.01766

33. Liu, S., Zhang, Y., Feng, Q., Qin, L., Pan, C., Lamin-Samu, A. T., & Lu, G. (2018). Tomato AUXIN RESPONSE FACTOR 5 regulates fruit set and development via the mediation of auxin and gibberellin signaling. Scientific reports, 8(1), 2971.

34. Livak, K. J., & Schmittgen, T. D. (2001). Analysis of relative gene expression data using real-time quantitative PCR and the 2− ΔΔCT method. methods, 25(4), 402–408.

35. Løvdal, T., & Lillo, C. (2009). Reference gene selection for quantitative real-time PCR normalization in tomato subjected to nitrogen, cold, and light stress. Analytical biochemistry, 387(2), 238–242.

36. Ma, Y. B., Li, W. L., Li, J. H., Li, M. J., Li, X. Y., Wei, C. M., … & Ma, X. R. (2025). Titanium ions promote tomato growth and increase stress resistance. BMC plant biology, 25(1), 995.

37. Maldonado, R. A., Bianchi, J. S., Chiesa, M. A., & Rodríguez, E. (2026). Streptomyces spp. alleviates drought stress and reduces yield losses by enhancing root development, net photosynthesis, and water-use efficiency in soybean plants. Plant Science, 113123.

38. Mastoraki, M., Bengoa Luoni, S., Lindeboom, J., Heuvelink, E., & Marcelis, L. F. (2025). Regulation of tomato fruit growth and development during different plant developmental stages by far red light: a combined anatomical, physiological, and transcriptomic analysis. The Plant Journal, 124(5), e70613.

39. Matsuo, S., Kikuchi, K., Fukuda, M., Honda, I., & Imanishi, S. (2012). Roles and regulation of cytokinins in tomato fruit development. Journal of experimental botany, 63(15), 5569–5579.

40. Modiba, M. P., Bell, T., & Babalola, O. O. (2025). Plant Growth Promoting Rhizobacteria and Bacterial Biocontrol Agents in Tomato Disease Management: Mechanisms, Applications, and Omics Perspectives. Global Challenges, 9(12), e00320.

41. Mesquita, A., Cerqueira, D., Rocha, M., Silva, D., Martins, C., & Souza, B. (2025). A review on rare and symbiotic actinobacteria: emerging biotechnological tools against antimicrobial resistance. Journal of Basic Microbiology, 65(6), e70036.

42. Ogugua, U. V., Ogbuewu, I. P., Mbajiorgu, C. A., & Adriaanse, P. (2025). Meta-analysis of biofertilizer effects of Bacillus species on tomato yield. Scientific Reports, 15(1), 34007.

43. Ozores-Hampton, M., Di Gioia, F., Sato, S., Simonne, E., & Morgan, K. (2015). Effects of nitrogen rates on nitrogen, phosphorous, and potassium partitioning, accumulation, and use efficiency in seepage-irrigated fresh market tomatoes. HortScience, 50(11), 1636–1643.

44. Porcel, R., Zamarreño, Á. M., García-Mina, J. M., & Aroca, R. (2014). Involvement of plant endogenous ABA in Bacillus megaterium PGPR activity in tomato plants. BMC plant biology, 14(1), 36.

45. Rafiq, M.; Guo, M.; Shoaib, A.; Yang, J.; Fan, S.; Xiao, H.; Chen, K.; Xie, Z. & Cheng, C. (2025) Unraveling the Hormonal and Molecular Mechanisms Shaping Fruit Morphology in Plants. Plants, 14, 974. 10.3390/plants14060974

46. R Core Team. (2025). R: A language and environment for statistical computing (Version 4.5.1) [Computer software]. Vienna, Austria: R Foundation for Statistical Computing.

47. Rehan, M., Al-Turki, A., Abdelmageed, A. H., Abdelhameid, N. M., & Omar, A. F. (2023). Performance of plant-growth-promoting rhizobacteria (PGPR) isolated from sandy soil on growth of tomato (Solanum lycopersicum L.). Plants, 12(8), 1588.

48. Renaudin, J. P., Cheniclet, C., Rouyère, V., Chevalier, C., & Frangne, N. (2023). The cell pattern of tomato fruit pericarp is quantitatively and differentially regulated by the level of gibberellin in four cultivars. Journal of Plant Growth Regulation, 42(9), 5945–5958.

49. Rodríguez, G., Pratta, G., Zorzoli, R., & Picardi, L. A. (2005). Caracterización de la generación segregante de un híbrido de tomate con genes nor y silvestres. Pesquisa Agropecuária Brasileira, 40(1), 41–46.

50. Rodríguez, G. R., Muños, S., Anderson, C., Sim, S. C., Michel, A., Causse, M., … & van Der Knaap, E. (2011). Distribution of SUN, OVATE, LC, and FAS in the tomato germplasm and the relationship to fruit shape diversity. Plant physiology, 156(1), 275–285.

51. Samaras, A., Roumeliotis, E., Ntasiou, P., & Karaoglanidis, G. (2021). Bacillus subtilis MBI600 promotes growth of tomato plants and induces systemic resistance contributing to the control of soilborne pathogens. Plants, 10(6), 1113.

52. Sayyed, R. Z., & Ilyas, N. (2024). as Biofertilizers, Biopesticides or Biostimulants for Improving the Crop. Plant Holobiome Engineering for Climate-Smart Agriculture, 71.

53. Schmülling, T., Werner, T., Riefler, M., Krupková, E., & Bartrina y Manns, I. (2003). Structure and function of cytokinin oxidase/dehydrogenase genes of maize, rice, Arabidopsis and other species. Journal of plant research, 116(3), 241–252.

54. Serrani, J. C., Sanjuán, R., Ruiz-Rivero, O., Fos, M., & García-Martínez, J. L. (2007). Gibberellin regulation of fruit set and growth in tomato. Plant physiology, 145(1), 246–257.

55. Shao, R. X., Xin, L. F., Zheng, H. F., Li, L. L., Ran, W. L., Mao, J., & Yang, Q. H. (2016). Changes in chloroplast ultrastructure in leaves of drought-stressed maize inbred lines. Photosynthetica, 54(1), 74–80.

56. Sundaresan, S., Philosoph-Hadas, S., Riov, J., Mugasimangalam, R., Kuravadi, N. A., Kochanek, B., … & Meir, S. (2016). De novo transcriptome sequencing and development of abscission zone-specific microarray as a new molecular tool for analysis of tomato organ abscission. Frontiers in plant science, 6, 1258.

57. Susic, Z., Pavlovic, N., Cvikic, D., & Sretenovic-Rajicic, T. (2000, October). Studies of correlation between yield and fruit characteristics of (Lycopersicon esculentum Mill.) hybrids and their parental genotypes. In II Balkan Symposium on Vegetables and Potatoes 579 (pp. 163–166).

58. Taheri, F., Jafar Daghighi, A., Bodaghi, H., Ebrahimi, A., & Ghasimi Hagh, Z. (2022). The effect of 1-methylcyclopropene application incorporated with packaging on the expression of ethylene-related genes and quality maintenance of cherry tomato cultivars. Research and Innovation in Food Science and Technology, 11(3), 275–290.

59. Vargas Beltrán, L. V. (2025). Efecto de Bacillus y Pseudomonas en la germinación y el crecimiento vegetal de Physalis peruviana.

60. Vurukonda, S. S. K. P., Giovanardi, D., & Stefani, E. (2018). Plant growth promoting and biocontrol activity of Streptomyces spp. as endophytes. International journal of molecular sciences, 19(4), 952.

61. Villafañe, D. L., Maldonado, R. A., Bianchi, J. S., Kurth, D., Gramajo, H., Chiesa, M. A., & Rodríguez, E. (2024). Streptomyces N2A, an endophytic actinobacteria that promotes soybean growth and increases yield and seed quality under field conditions. Plant science, 343, 112073.

62. Villafañe, D. L., Maldonado, R. A., Rodríguez, E., & Chiesa, M. A. (2025). Endophytic Streptomyces sp. N2A protects soybean against fungal diseases through two distinct mechanisms: David L. Villafañe et al. BioControl, 70(4), 529–542.

63. Wang, R., & de Maagd, R. A. (2025). Transcriptional control of tomato fruit development and ripening. Journal of Experimental Botany, 76(21), 6311–6326.

64. Wang, J., Gao, X., Yang, M., Zhang, Y., & Chen, Y. (2025). Alleviative effects of plant growth-promoting rhizobacteria on salt-stressed rice seedlings: mechanisms mediated by rhizosphere microbiota and root exudates. Frontiers in Plant Science, 16, 1661074.

65. Yan, N., Wang, W., Mi, T., Zhang, X., Li, X., & Du, G. (2024). Enhancing tomato growth and soil fertility under salinity stress using halotolerant plant growth-promoting rhizobacteria. Plant Stress, 14, 100638.

66. Yang, P., Condrich, A., Scranton, S., Hebner, C., Lu, L., & Ali, M. A. (2024). Utilizing plant growth-promoting rhizobacteria (PGPR) to advance sustainable agriculture. Bacteria, 3(4), 434–451.

67. Zhang, S., Xu, M., Qiu, Z., Wang, K., Du, Y., Gu, L., & Cui, X. (2016). Spatiotemporal transcriptome provides insights into early fruit development of tomato (Solanum lycopersicum). Scientific Reports, 6(1), 23173.

68. Zhang, Y., Zhao, S. Y., Zhang, R. H., Li, B. L., Li, Y. Y., Han, H., … & Chen, Z. J. (2024). Screening of plant growth-promoting rhizobacteria helps alleviate the joint toxicity of PVC+ Cd pollution in sorghum plants. Environmental Pollution, 355, 124201.

